# BreakCA, a method to discover indels using ChIP-seq and ATAC-seq reads, finds recurrent indels in regulatory regions of neuroblastoma genomes

**DOI:** 10.1101/605642

**Authors:** Arko Sen, Sélène T. Tyndale, Yi Fu, Galina Erikson, Graham McVicker

**Author notes:** Corresponding Authors: AS, GM.

## Abstract

Most known cancer driver mutations are within protein coding regions of the genome, however, there are several important examples of oncogenic non-coding regulatory mutations. We developed a method to identify insertions and deletions (indels) in regulatory regions using aligned reads from chromatin immunoprecipitation followed by sequencing (ChIP-seq) or the assay for transposase-accessible chromatin (ATAC-seq). Our method, which we call BreakCA for Breaks in Chromatin Accessible regions, allows non-coding indels to be discovered in the absence of whole genome sequencing data, out-performs popular variant callers such as the GATK-HaplotypeCaller and VarScan2, and detects known oncogenic regulatory mutations in T-cell acute lymphoblastic leukemia cell lines. We apply BreakCA to identify indels in H3K27ac ChIP-seq peaks in 23 neuroblastoma cell lines and, after removing common germline variants, we identify 23 rare germline or somatic indels that occur in multiple neuroblastoma cell lines. Among them, 4 indels are candidate oncogenic drivers that are present in 4 or 5 cell lines, absent from the genome aggregation database of over 15,000 whole genome sequences, and within the promoters or first introns of known genes (*PHF21A, ADAMTS19, GPR85* and *RALGDS*). In addition, we observe a rare 7bp germline deletion in two cell lines, which is associated with high expression of the histone demethylase *KDM5B*. Overexpression of *KDM5B* is prognostic for many cancers and further characterization of this indel as a potential oncogenic risk factor is therefore warranted.

## Introduction

Several non-coding mutations are known to be important oncogenic drivers. For example, mutations within the promoter of *TERT* are extremely common and cause its overexpression in numerous cancers[1–3] and 2-12bp insertions create new enhancer sequences that drive overexpression of *TAL1* in 4-6% of T-cell acute lymphoblastic leukemias (T-ALLs)[4]. Genome-wide scans for recurrent non-coding mutations have found a handful of additional candidates including recurrent mutations in regulatory regions upstream of *PLEKHS1, WDR74*, and *SDHD*[5]. A somatic mutation screen using WGS from chronic lymphocytic leukemia patients identified recurrent T>C mutations in the 3’UTR of *NOTCH1* which cause it to be aberrantly spliced, as well as clustered mutations across multiple patients in an enhancer region for PAX5[6]. Non-coding variants can also reposition regulatory sequences so that they activate oncogenes[4, 7, 8] and some non-coding germline polymorphisms that disrupt factor binding motifs are associated with cancer risk. A single-nucleotide polymorphism (SNP) upstream of *MYC* impacts binding of the YY1 transcription factor and is associated prostate cancer[9], SNPs in OCT1/RUNX2 and C/EBPβ binding sites near *FGFR2* modulate its expression and are associated with breast cancer[10], and a SNP that disrupts a GATA3 binding site in an *LMO1* enhancer is associated with neuroblastoma[11].

While the cost of whole-genome sequencing (WGS) has decreased dramatically, it remains expensive for large panels of individuals and functional interpretation of non-coding mutations is difficult. One way to overcome these challenges is to identify genetic variants using data from experiments such as chromatin immunoprecipitation followed by sequencing (ChIP-seq) and the Assay for Transposase-Accessible Chromatin (ATAC-seq). These experiments generate sequence reads from regulatory regions of the genome, which can potentially be used to identify non-coding driver mutations in cancer samples. Here we describe a new method to identify indels from ChIP-seq and ATAC-seq reads, which we call BreakCA, for “Breaks in Chromatin Accessible” regions. We assess the performance of BreakCA on ATAC-seq and ChIP-seq reads from the GM12878 lymphoblastoid cell line and the Jurkat T-ALL cell line and verify that BreakCA detects known oncogenic indels that create enhancers for the *TAL1* and *LMO2* genes[4, 12]. We then apply BreakCA to H3K27ac ChIP-seq data from 23 neuroblastoma cell lines. After filtering the indels using a large database of known germline variants, we identify recurrent rare germline or somatic indels that may be oncogenic drivers for neuroblastoma.

## Results

### Detecting indels with BreakCA

We hypothesized that aligned sequences from ChIP-seq and ATAC-seq experiments can be used to identify indels in regulatory regions of the genome in the absence of whole-genome sequencing (WGS). To test this hypothesis, we developed a method to detect indels by exploiting properties of mapped reads such as gaps in alignments (i.e. insertions or deletions) and clipping at read ends (Fig 1a). Our method, which we call BreakCA for “Breaks in Chromatin Accessible” regions, collects 16 features from mapped reads and uses a random forest to identify 20bp windows that contain indels.

**Figure 1:**
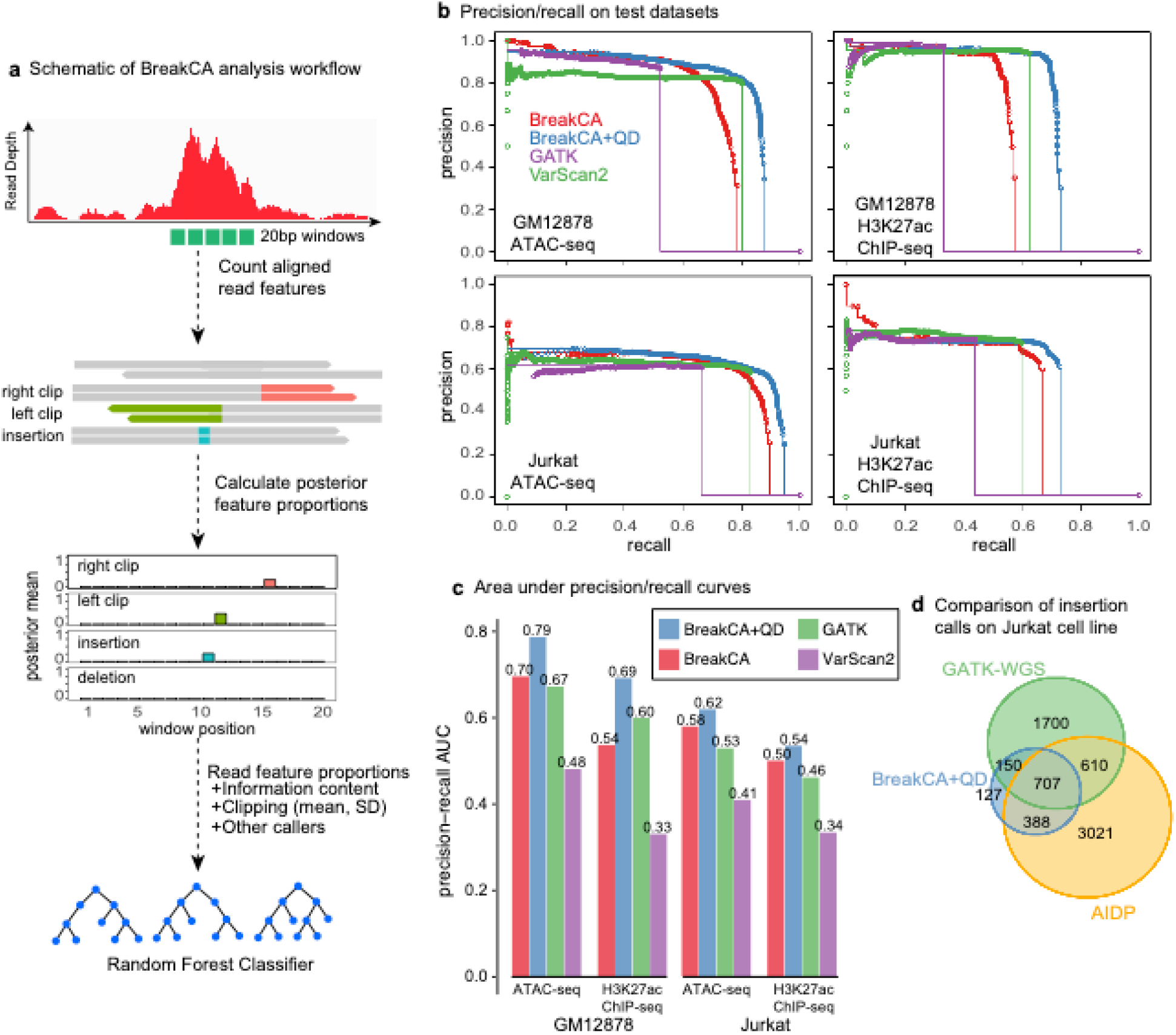
**a)** BreakCA detects insertions and deletions (indels) in cancer genomes using mapped ATAC-seq or ChIP-seq reads. **b)** Precision-recall curves for indels scored by four callers: BreakCA, BreakCA+QD (BreakCA with GATK Quality Depth), the GATK-HaplotypeCaller and VarScan2. Each panel shows precision recall curves for a different test dataset: 50bp paired-end ATAC-seq from GM12878 (2980 true and 842,738 false windows); 50bp single-end H3K27ac ChIP-seq from GM12878 (623 true and 254,328 false windows); 42bp paired-end ATAC-seq from Jurkat (5020 true and 1,023,159 false windows) and 40bp single-end H3K27ac from Jurkat (6403 true and 1,267,161 false windows). Indels called by Platinum Genomes are used as the “ground truth” for GM12878 and indels called by the GATK-HaplotypeCaller applied to whole-genome sequence (WGS) data are used as the “ground truth” for Jurkat. **c)** Area under the precision recall curves for each of the test datasets and indel callers. **d)** Comparison of insertions identified in the Jurkat cell line from: GATK-HaplotypeCaller applied to WGS and then filtered for ChIP-seq peaks; BreakCA applied to H3K27ac ChIP-seq; Abraham’s Insertion Detection Pipeline (AIDP) applied to H3K27ac ChIP-seq.

To train BreakCA and assess its performance, we created separate training and test datasets from 50bp paired-end ATAC-seq data and 50bp single-end H3K27ac ChIP-seq data from the GM12878 lymphoblastoid cell line[13, 14]. To label testable windows within ChIP-seq and ATAC-seq peaks as “true” or “false” we used indel calls from the Platinum Genomes (PG) project as known positives[15]. After training the random forest on the GM12878 training dataset, we evaluated its performance on the test dataset and compared it to two popular variant callers: VarScan2[16] and the GATK-HaplotypeCaller[17].

We quantified overall performance using the area under precision-recall curves (Fig 1b,c) and found that BreakCA (prAUC=0.70) performs better than VarScan2 (prAUC=0.48) and comparably to the GATK-HaplotypeCaller (prAUC=0.67) for paired-end ATAC-seq data. For single-end ChIP-seq data BreakCA (prAUC=0.54) performs better than VarScan2 (prAUC=0.33) but worse than the GATK-HaplotypeCaller (prAUC=0.60). An important advantage of BreakCA is that additional features, including output from other variant callers, can be easily added to improve its performance. We added the Quality Depth (QD) reported by the GATK-HaplotypeCaller as a feature for BreakCA and observed substantial improvements in the prAUC for both the ChIP-seq (prAUC=0.69) and ATAC-seq (prAUC=0.79) datasets such that its performance was substantially better than both GATK and VarScan2. We call this version of our method BreakCA+QD.

To test the performance of BreakCA on a cancer cell line that was not used for training, we used paired-end ATAC-seq and H3K27ac ChIP-seq data from the Jurkat T-ALL cell line and obtained WGS data for the same cell line from a published study[18]. Since there is no gold-standard set of indel calls for this cell line, we used indels identified by the GATK-HaplotypeCaller run on WGS data as our “ground truth”. The performance of BreakCA (prAUC= 0.58) on the Jurkat ATAC-seq data was better than both VarScan2 (prAUC=0.41) and GATK-HaplotypeCaller (prAUC=0.53) and improves further when information from GATK is included (prAUC=0.62). For single-end ChIP-seq, while BreakCA (prAUC=0.50) out-performed VarScan2 (prAUC=0.34), its overall performance was comparable to GATK-HaplotypeCaller (prAUC= 0.46) and we observed an improvement in prAUC after adding QD from GATK (prAUC=0.54). While all methods appear to perform worse on the Jurkat datasets compared to the GM12878 datasets (Fig 1c), it is important to note that the performance is greatly underestimated due to inaccuracies in the Jurkat “ground truth” dataset (high false-negative rates) compared to the high-quality platinum genomes dataset.

We compared BreakCA to an orthogonal method that was recently developed by Abraham et al. to identify small insertions from ChIP-seq reads[12]. This method, which we refer to as Abraham’s Insertion Detection Pipeline (AIDP), assembles contigs from ChIP-seq reads that fail to map the reference genome. AIDP was previously applied to H3K27ac ChIP-seq data from the Jurkat Cell line and we ran BreakCA on the same dataset. We compared the insertion calls from AIDP and BreakCA to those from the GATK-HaplotypeCaller, which was run on WGS data (Fig 1f). While BreakCA detects a much smaller number of insertions (n=1372 for BreakCA compared to n=4726 for AIDP), BreakCA’s overlap with the WGS-identified indels is far higher (62% for BreakCA compared to 28% for AIDP). Of the 515 BreakCA insertions that are not detected by GATK, most (75%) are also detected by AIDP. Only 9% of BreakCA-identified indels are not called by either GATK or AIDP, suggesting that the accuracy of BreakCA is high. In contrast, most of the insertions detected by AIDP (64%), are only detected by AIDP, suggesting that a large proportion of them may be false-positives.

Our performance evaluations on GM12878 and Jurkat T-cells indicate that BreakCA+QD offers the best balance of precision and recall for both ATAC-seq and ChIP-seq datasets and we used this approach for all subsequent analyses. We selected score thresholds based on the precision-recall curves for the GM12878 dataset. Specifically, we used a score threshold of ≥0.60 corresponding to ~86% precision and ~75% recall for paired-end ATAC-seq and a score threshold of ≥0.33 corresponding to ~0.91% precision and ~69% recall for single-end ChIP-seq data.

To test whether BreakCA detects known oncogenic mutations, we applied it to ATAC-seq from four T-ALL cell lines (Jurkat, MOLT-4, CCRF-CEM and RPMI-8402) and one CML cell line (K-562). Two of these cell lines (Jurkat and MOLT-4) are known to harbor oncogenic insertions 8kb upstream of the TAL1 promoter[4] and BreakCA successfully detects both of them (Fig 2). In addition, we verify that BreakCA detects an insertion that is known to be associated with allele-specific expression of the *LMO2* oncogene in MOLT-4 cells[12] (Supplementary Fig 1). These results indicate that BreakCA can detect oncogenic indels in regulatory regions of the genome using ATAC-seq or ChIP-seq reads. We next applied BreakCA to characterize the non-coding regulatory landscape of neuroblastoma.

**Figure 2:**
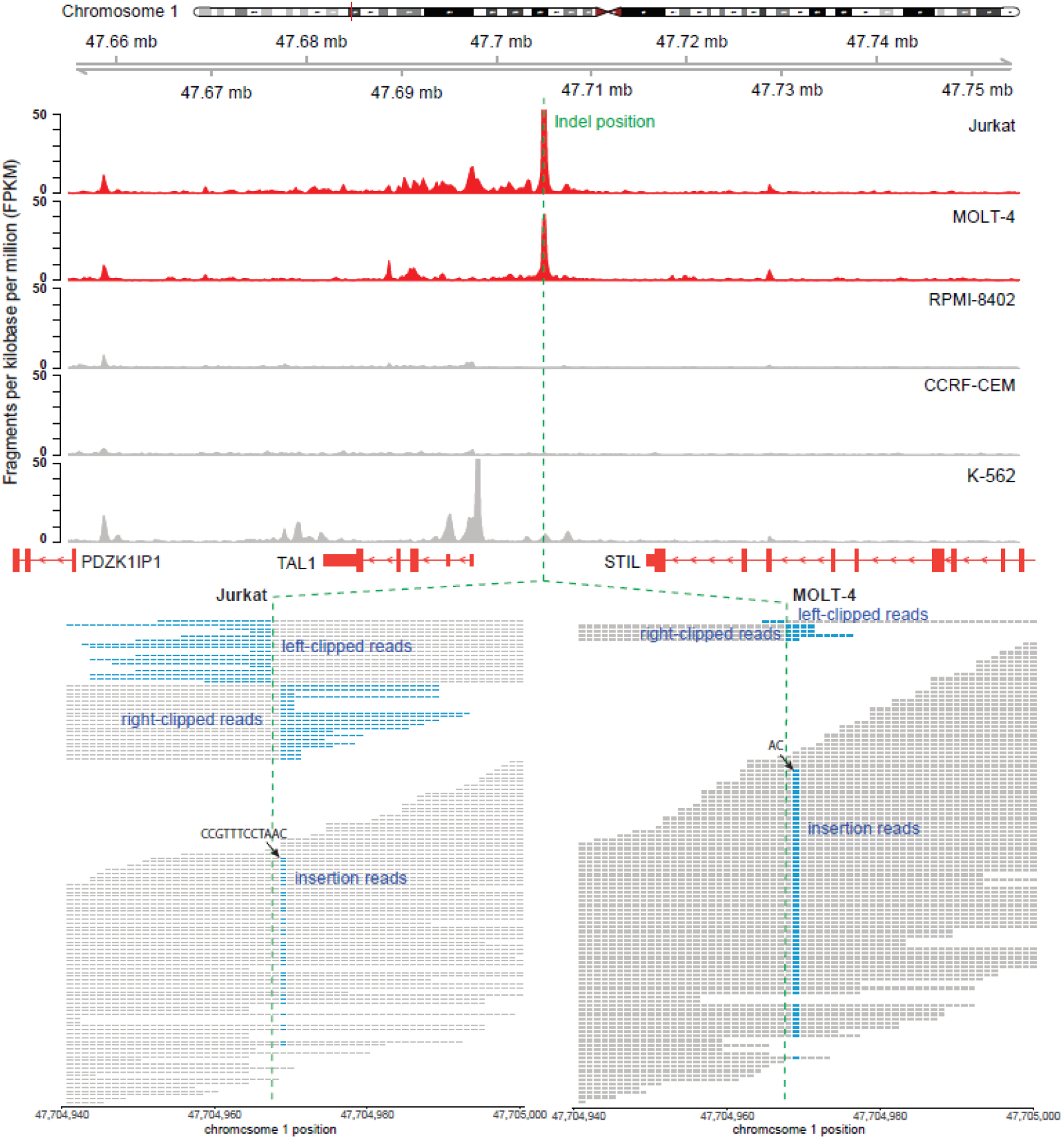
BreakCA detects known oncogenic insertions upstream of *TAL1* in the Jurkat and MOLT-4 cell lines. The insertion is well-covered by both clipped and insertion-containing reads and inspection of the read pileup reveals a *CCGTTTCCTAAC* insertion in Jurkat and an *AC* insertion in MOLT-4 cell lines (bottom panel). Mapped reads per base position are in grey and soft-clipped bases and insertion positions are colored in blue.

### Discovery of indels in neuroblastoma cell lines

Neuroblastoma (NB) is a childhood cancer of the peripheral nervous system with low mutation rates and few recurrently mutated genes[19–22]. Many neuroblastoma tumors harbor no known oncogenic mutations and it has been hypothesized that many high-risk neuroblastomas are driven by rare germline variants, copy number alterations or epigenetic modifications that occur during tumor evolution[19]. We hypothesized that recurrent indels in regulatory regions may also be important drivers of NB. To test this hypothesis, we obtained H3K27ac ChIP-seq from 26 NB cell lines and 2 normal human neural crest cells (hNCCs)[23] and ran BreakCA on these samples to identify indels within regulatory regions defined by H3K27ac peaks.

The SH-EP and SH-SY5Y cell lines are subclones of the SK-N-SH cell line, so we assigned variant windows identified in these lines to SK-N-SH and treated the combined set of indels as a single cell line. We noticed that the genotypes for the GICAN cell line are nearly identical to those of the GIMEN cell line and that the *SRY* expression of the GICAN cell line is inconsistent with the reported sex (annotated male with no *SRY* expression). We concluded that the GICAN cell line is probably mislabeled and excluded it from further analyses. Our final set of analyzed cell lines consisted of 23 NB cell lines and 2 hNCCs.

We implemented a filtering pipeline to remove potential artefacts and common germline indels (Fig 3a). First, we removed windows overlapping regions that were previously-identified as problematic for ChIP-seq analysis based on their high ratio of multi-mapping to unique mapping reads[24]. Second, we removed indel-containing windows that were detected in the two hNCCs as these are likely to be common germline variants. Third, we conservatively removed 16,234 windows that contained short tandem repeat (STR) sequences (Supplementary Fig 2). While STRs have very high mutation rates and contain many true indels, they are also prone to a high rate of variant-calling artifacts caused by polymerase slippage during PCR[25–27].

**Figure 3:**
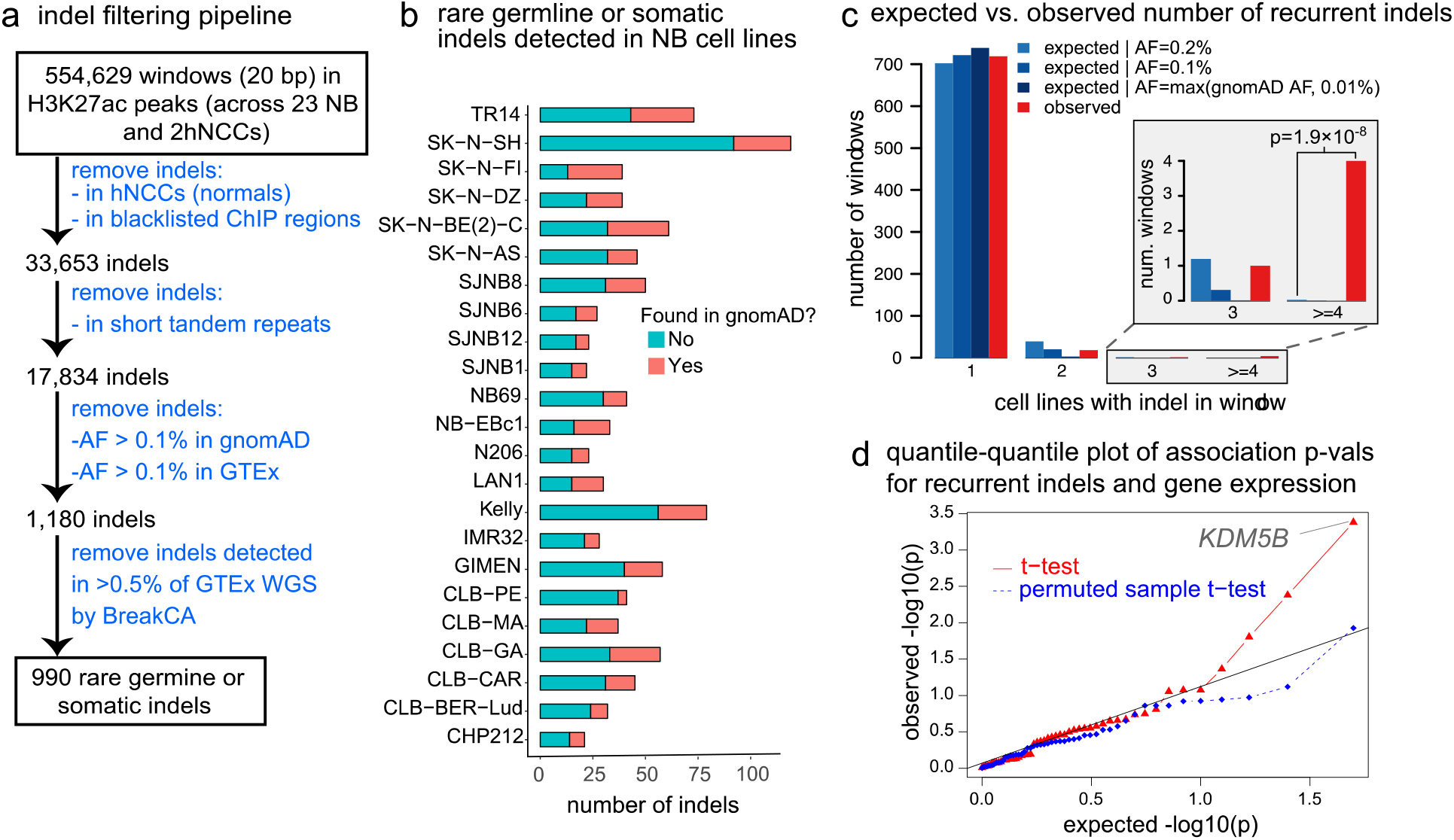
**a)** Filtering pipeline for identifying rare germline or somatic (RS) indels in 23 neuroblastoma cell-lines. **b)** The number of rare germline or somatic indel windows detected in each neuroblastoma cell-line after filtering, divided into those that are present/absent in gnomAD+GTEx **c)** The expected number of windows containing RS indels in one or more cell lines (blue bars), assuming that they are inherited germline variants with given allele frequencies (AFs). The plots are conditional on seeing the indel in at least one cell line, and on the number of testable cell lines being at least 5. The red bars are the observed number of windows with one or more RS indels. **d)** Quantile-quantile plot of expected and observed - log10 p-values for RS indel-gene pairs. Each RS indel was tested for association with the expression of all genes within 100kb.

Since there are no matched normal tissues for the NB cell lines, germline variants cannot be distinguished from somatic mutations. To eliminate common germline indels and focus on those that are either rare germline variants or somatic mutations, we filtered indels based on their allele frequency in samples from the Genotype-Tissue Expression Project (GTEx) and the genome aggregation database (gnomAD) of 15,708 whole genome sequences[28, 29]. We found 1,180 out of 17,834 windows contained indels that were completely absent from GTEx and gnomAD or present with an allele frequency of less than 0.1% (Fig 3a). Additionally, we ran BreakCA on WGS data from 300 GTEx samples and removed indels that we detected in greater than 0.5% of samples. The 990 indel windows that remained after these filtering steps contain either rare germline or somatic indels, which we refer to as RS indels.

### Recurrent rare germline or somatic indels

RS indels that occur in multiple cell lines are more likely to be oncogenic drivers. To ask how many of the 20bp windows contained recurrent RS indels, we focused on the 742 windows that contained at least one such indel and that were testable by BreakCA (i.e. windows with at least 10 ChIP-seq reads) in 5 or more cell lines. In total, 23 windows contained RS indels in two or more NB cell lines. Remarkably, 4 windows contained an RS indel that is present in 4 or 5 cell lines. RS indels are very unlikely to occur in 4 or more cell lines due to random inheritance of rare genetic variants (P < 1.8×10^−9^ by Poisson test; Fig 3c) and therefore these indels may be highly-recurrent oncogenic driver mutations. Recurrent indels can also arise due to high mutation rates, however this scenario is unlikely given that (1) the total number of RS indels that we observe in each cell line is low (Fig 3b) and (2) we have removed windows that overlap annotated STRs, which are typically the most mutagenic sequences. Recurrent rare variants could also be observed if some of the cell lines were derived from close relatives, however, none of the cell lines appear to be closely related because the total number of RS indels shared between them is low, and the 4 highly-recurrent RS indels are present in different subsets of cells (Table 1).

**Table 1:** Rare germline or somatic (RS) indels that are observed in at least 4 cell lines.

| Window | Testable cell lines | Cell lines with indel | gnomAD AF | Gene | Location | Alleles | Notes |
| --- | --- | --- | --- | --- | --- | --- | --- |
| chr11:<br>46,144,281-46,144,300 | 21 | CLB-GA,<br>SJNB12,<br>SJNB6,<br>SK-N-BE (2)-C | 0 | PHF21A | Promoter | C/CG | Insertion in all cell-lines |
| chr5:<br>128,796,561-128,796,580 | 16 | CLB-CAR,<br>CLB-GA,<br>CLB-PE,<br>SJNB6 | 0 | ADAMTS19 | First Intron | T/TC (CLB-CAR, CLB-GA, CLB-PE),<br>C/CT (SJNB6) | Insertion in all cell lines but insertion position differs |
| chr7:<br>112,726,881-112,726,900 | 19 | CLB-GA,<br>CLB-PE,<br>SJNB6,<br>SK-N-AS,<br>TR14 | 0 | GPR85 | First Intron | Complex event | 1bp insertion and multiple mismatches to reference |
| chr9:<br>135,994,681-135,994,700 | 22 | CLB-PE,<br>SJNB1,<br>SK-N-AS,<br>TR14 | 0 | RALGDS | First Intron | T/TC | Insertion in all cell lines |

All of the RS indels in the highly recurrent windows (i.e. windows containing RS indels in at least 4 cell lines) are not observed in gnomAD, and are close to transcription start sites (TSSs) of protein-coding genes (Table 1). The first window contains a 1bp insertion that is 295bp upstream of the TSS of *PHF21A* and is observed in 4 cell lines. The second window is in the first intron of *RALGDS* and contains an insertion that is present in 4 cell lines. The third window is within the first intron of *ADAMTS19* and contains 2 insertions that are 5bp apart: a C insertion (present in 3 cell lines) and a T insertion (present in 1 cell line). Finally, the fourth window is within the first intron of *GPR85* and contains a complex event consisting of a 1bp insertion and multiple mismatches to the reference sequence (Supplementary Fig 5). Since this complex event appears to be the same in the 5 cell lines where it is detected it may be a rare germline haplotype.

### An intronic germline deletion associated with *KDM5B* expression

We hypothesized that recurrent RS indels might be associated with the expression of nearby genes and we therefore tested all genes located within 100kb of RS indels for differences in expression using Student’s t-test (assuming equal variances). The p-values from these tests show a clear departure from the null expectation (Fig 3d), however our power to detect associations is limited by the fact that each indel is only present in 2-5 cell lines. Under a stringent false-discovery rate threshold of 5%, a single test, between an indel and the expression of the H3K4me3/me2 Lysine Demethylase 5B (*KDM5B*), is significant (nominal p-value=4.1×10”^4^; Benjamini-Hochberg adjusted p-value=0.021).

The indel associated with *KDM5B* expression is a 7bp deletion (*GCCTCGG*/-), which is located in its first intron and is present only in the SJNB1 and NB-EBc1 cell lines (Fig 4a & Supplementary Fig 4). This deletion is a germline variant that occurs at a very low minor allele frequency in both gnomAD (1.3×10^−4^) and GTEx (7.9×10^−4^). The expression of *KDM5B* is very high in the SJNB1 and NB-EBc1 cell lines compared to cell lines that do not contain the indel (with exception of SJNB12) as well as tissues that may resemble the cell-type of origin for NB including hNCCs and adrenal and spinal tissues from GTEx (Fig 4b). We performed a motif analysis (see methods) for the 40bp region centered on the indel and found binding motifs for Transcription Factor AP-2 Beta (TFAP2B) and Gamma (TFAP2C) in the reference sequence, which co-localize with the deletion. The indel disrupts the TFAP2B/2C motif and creates a ZNF263 motif in its place (Fig 4c). *TFAP2B* is highly expressed in many NB cell lines (including SJNB1 and NB-EBc1), whereas *ZNF263* appears to be expressed across both normal hNCC and NB cells (Supplementary Fig 3).

**Figure 4:**
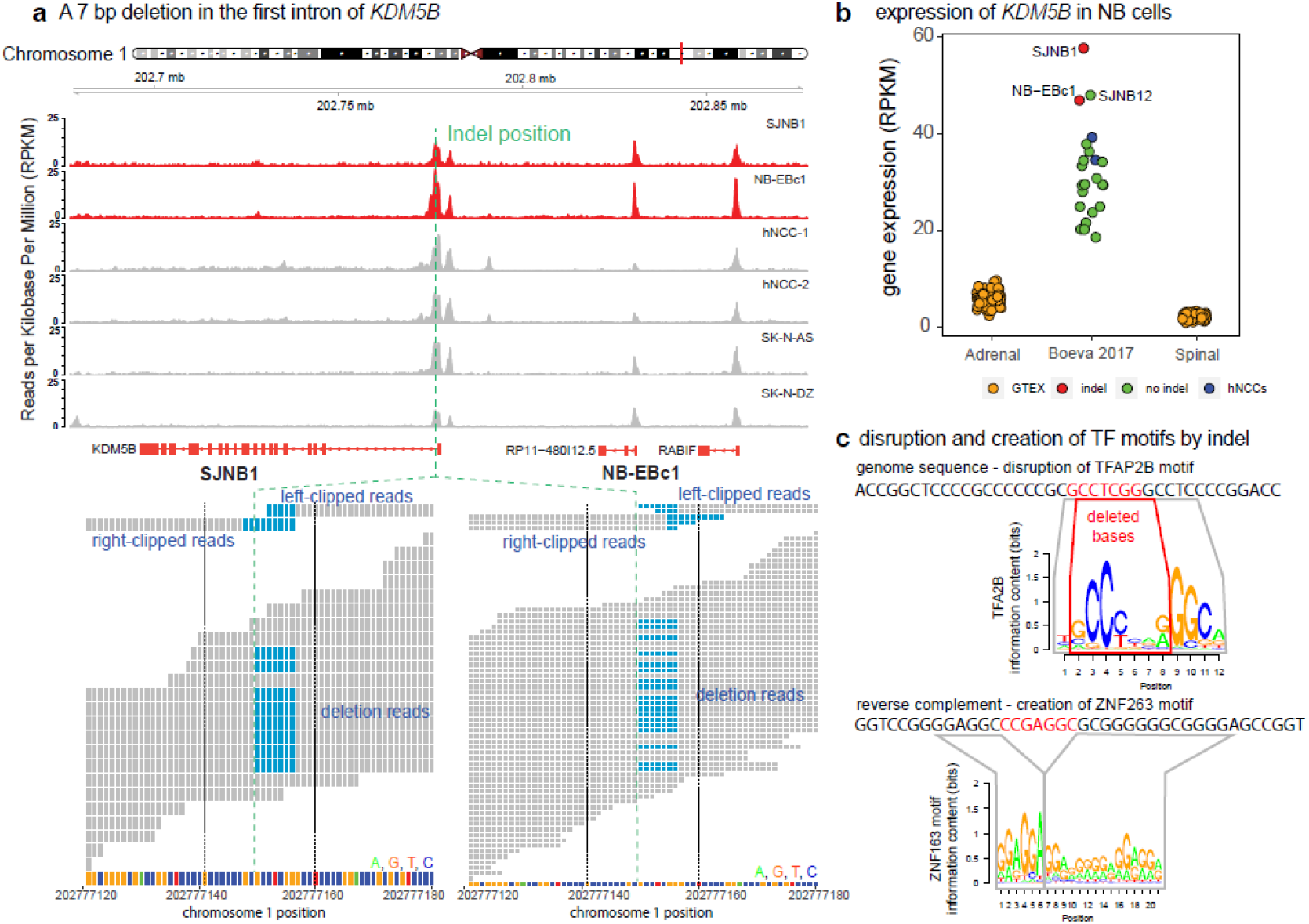
**a)** H3K27ac read depth for two indel-containing neuroblastoma cell lines (in red), 2 human neural crest cell lines, and two non-indel neuroblastoma cell lines. The germline deletion is located within the first intron of *KDM5B* and is covered by both soft-clipped and deletion reads in both cell lines where it is detected. Soft-clipped and deletion base positions are colored in blue. **b)** *KDM5B* gene expression in Reads Per Kilobase Per Million mapped reads (RPKM) in neuroblastoma (NB) cell-lines (from Boeva et al. 2017), human neural crest cells (hNCCs), and in adrenal and spinal tissues from the GTEx project **c)** The *GCCTCGG/*- 7bp deletion disrupts a TFAP2B motif (top panel) and creates a new ZNF263 motif.

## Discussion

We used BreakCA to identify indels in 23 neuroblastoma cell lines. One of the most interesting events we detected is a 7bp deletion which replaces a *TFAP2B* binding motif with a *ZNF263* motif within the first intron of *KDM5B* in the SJNB1 and NB-EBc1 cell lines (Fig 4). This indel appears to be germline, as an identical event is detected in 4 out of 31,266 chromosomes surveyed by gnomAD. The presence of this deletion is not associated with a difference in H3K27ac levels but is associated with overexpression of *KDM5B* in these cells, most likely through disruption of the TFAP2B motif. *KDM5B* has known functions in NB and its knockdown results in a 5-fold decrease in cell motility and suppresses the epithelial-mesenchymal transition via downregulation of *NOTCH1* expression in NB cells[30]. Furthermore, NB cells that overexpress *KDM5B* form spheroids that are more resistant to in vitro treatment with doxorubicin, etoposide and cisplatin[30]. Finally, *KDM5B* overexpression is associated with poor outcomes in several other cancers including glioma, hepatocellular carcinoma, non-small cell lung cancer, and prostate cancer[31–34].

The disruption of the TFAP2B motif is also interesting because the *TFAP2B* transcription factor is highly expressed in migrating neural crest cells which are involved in the development of the sympathetic nervous system and are likely cells of origin for NB[35]. In addition, low *TFAP2B* expression is associated with poor survival and prevents neuronal differentiation of NB cells in vitro via downregulation of *MYCN* and REST[36]. Our results indicate that TFAP2B may also downregulate *KDM5B*.

The recurrent RS indels that are present in 4 or 5 cell lines may be oncogenic mutations or rare germline predisposition variants. One of the indels is upstream of *PHF21A*, which encodes a subunit of the BRAF-histone deacetylase complex that is recruited by REST to silence neuronal-specific genes[37]. Another indel is in the first intron of *RALGDS*, which is a guanine exchange factor in a Ras signaling pathway[38]. REST is known to inhibit neuronal differentiation of neuroblastoma cells[39, 40] and members of Ras signaling pathways are frequently mutated in relapsed neuroblastoma[41], so both of these indels are excellent candidates for further functional characterization.

In conclusion, BreakCA allows ATAC-seq and ChIP-seq experiments to be treated as “exome capture” for the regulatory genome and enables the discovery of oncogenic regulatory indels in the absence of WGS data. While we focused only on short indels in this study, future studies could combine indels called by BreakCA with single nucleotide variants and larger events such as chromosome translocations and copy number alterations. A caveat of BreakCA is that it cannot detect variants outside of ChIP-seq or ATAC-seq peaks or variants that cause a complete loss of these peaks. However, despite this limitation, BreakCA paired with rigorous filtering of common germline events, can identify potential cancer driver mutations and germline risk variants that increase or at least maintain the regulatory activity of a sequence. As a proof-of-principal we used BreakCA to identify recurrent indels in NB cell lines that may be important for neuroblastoma progression, metastasis and drug resistance.

## Methods

### ATAC-seq and ChIP-seq data

ATAC-seq for the GM12878 lymphoblastoid cell was obtained from Buenrostro et al. 2013 (GEO: GSE47753)[13]. H3K27ac ChIP-seq for the GM12878 lymphoblastoid cell line was obtained from Ernst et al. 2011 (GEO: GSE26320)[14]. H3K27ac ChIP-seq and RNA-seq data from 26 Neuroblastoma (NB) cell lines was obtained from Boeva et al. 2017 (GEO: GSE90683)[23]. H3K27ac ChIP-seq (75bp single-end) for the Kelly NB cell line was obtained from Zeid et al. 2018 (GEO: GSE80151)[42]. ATAC-seq experiments for the Jurkat, CCRF-CEM, RPMI-8402 and MOLT-4 T-ALL cell lines and the K-562 chronic myelogenous leukemia cell line were performed in our lab.

### ATAC-seq experiments

ATAC-seq experiments for the Jurkat, MOLT-4, CCRF-CEM, RPMI-8402 and K-562 cell lines were performed using the Omni-ATAC-seq method as described[43], with minor modifications. In each experiment, 1×10^5^ cells were centrifuged at 1000 × g for 10 min at 4 °C. Following aspiration, a cell count of the supernatant was performed, the remaining cell number was calculated, and all further reagents in the protocol were titrated to this cell number. For every 5×10^4^ cells, nuclei were isolated in 50 μl cold ATAC-Resuspension Buffer (RSB) (10 mM Tris-HCl pH 7.4, 10 mM NaCl, 3 mM MgCl2) containing 0.1% NP40, 0.1% Tween-20, and 0.01% Digitonin, and pipet-mixed up-and-down at least 5 times. Nuclei isolation mix was incubated on ice for 3 exactly minutes, washed in 1 ml of cold ATAC-RSB containing 0.1% Tween-20 (but no NP40 or Digitonin) and centrifuged at 1000 × g for 10 min at 4 °C. Nuclear DNA was tagmented in 50 μl Transposition mix (25 μl 2 × TD buffer, 2.5 μl transposase (100nM final), 0.5 μl 1% digitonin, 0.5 μl 10% Tween-20, 16.5 μl PBS and 5 μl diH2O), and incubated in a thermomixer at 37 °C, 1000 × g for 30 min. Tagmented DNA was purified with Zymo DNA Clean and Concentrator-5 Kit (cat# D4014). Library amplification was performed using custom indexing Nextera primers from IDT in a 50 μl Kapa Hi Fi Hot Start PCR reaction (cat# KK2602). Following 3 initial cycles, 1 μl of PCR reaction was used in a quantitative PCR (Kapa qPCR Library Quantitation Kit cat# KK4824) to calculate the optimum number of final amplification cycles. Library amplification was followed by SPRI size selection with Kapa Pure Beads (cat# KK8002) to retain only fragments between 80-1,200bp. Library size was obtained on an Aglient Bio-Analyzer or TapeStation using a High Sensitivity DNA kit and factored into final Kapa qPCR results to calculate the final size-adjusted molarity of each library. Libraries were pooled and sequenced on an Illumina NextSeq500 in Paired-End 42 base pair configuration at the Salk Next Generation Sequencing Core. ATAC-seq data quality was assessed using the fraction of reads within peaks (FRiP) and fraction of mitochondrial reads (Fmito) metrics (Supplementary Table 1).

### Aligning reads to the genome and calling peaks

ChIP-seq and ATAC-seq reads were aligned to the hg19/GRCh37 reference genome using BWA-MEM (version 0.7.15-r1140)[44] with default parameters. BWA-MEM performs local alignment and retains reads that only partially map to the genome as soft-clipped alignments, which are useful for identifying indels. Reads were filtered using samtools (version=0.1.19)[45] and only non-duplicated reads with mapping quality (MAPQ ≥ 30) were kept for downstream analysis. ChIP-seq and ATAC-seq peak regions were identified using MACS2 (version 2.1.1)[46] with default parameters.

### RNA-seq and gene expression

RNA-seq reads were aligned to the hg19/GRCh37 reference genome using STAR (version STAR_2.5.3a). Mapped reads were filtered using samtools (version 0.1.19) and only non-duplicated reads with mapping quality (MAPQ ≥ 20) were kept for expression analysis. Read counts per gene were calculated with featureCounts (version 1.6.3)[47] using the GENCODE v19 GTF file in paired-end mode and converted to RPKM using edgeR[48].

### Obtaining features for indel prediction

We divided the genome into non-overlapping 20bp windows and collected 16 features from each window that could help predict the presence of an indel (Supplementary Table 2). Many of the features are based on the proportion of read alignments that contain characteristics such as insertions, deletions, or clipping. For example, one of the features that we consider is the proportion of reads that contain an insertion starting at one base position. We obtain posterior estimates of these proportions using an empirical Bayesian approach, which prevents over-estimation of the proportion when the number of aligned reads at a position is small (i.e. this is a Bayesian alternative to using a pseudocount). We assume that the count of reads with a given characteristic (e.g. insertion) at genomic position *i*, is a binomially-distributed random variable, *X_i_* with proportion parameter *p_i_:*

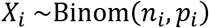

where, *n_i_* is the total number of reads overlapping genomic position *i*. We place a Beta prior on*p_i_*:

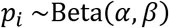

with *α* and *β* hyperparameters that describe the shape of the distribution. We estimate α and β empirically using the estimated proportion mean (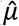) and variance (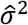) computed across all positions within peaks:

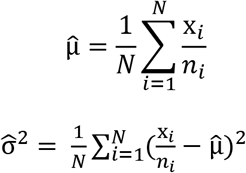

where *N* is the total number of positions, *n_i_* is the number of reads overlapping position *i*, and *x_i_* is the number of reads with a characteristic at that position (e.g. insertion). The *α* and *β* hyperparameters are then calculated as:

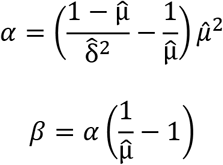

The Beta prior distribution is conjugate with the Binomial likelihood and the corresponding posterior distribution for the proportion is:

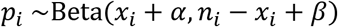

Using this distribution, we calculate the posterior mean and posterior standard deviation at each site, *i*, for the following proportions: (1) insertion reads, (2) deletion reads, (3) reads with clipping on the left side, (4) reads with clipping on the right side. We only perform posterior calculations for base positions covered by at least 10 reads.

For positions with left-clipped or right-clipped reads we collect additional clipping features including (1) the mean clipping length, (2) the standard deviation in clipping length, and (3) the information content of the clipped sequences (see below).

We compute the above features for each site, but we perform predictions on windows containing 20 sites. To assign features to 20bp windows we use the sites with the highest posterior means for each of the 4 proportion types. Additional features related to clipped reads, such as the mean right clipping length, are taken from the sites with the highest posterior mean clipping proportions.

### Information content of clipped reads

We compute an information content (*IC*) for clipped reads, which is defined so that highly similar clipped sequences have high *IC*, and dissimilar clipped sequences (perhaps arising from multiple genomic locations) have low *IC*. To compute the *IC* of overlapping clipped reads that start clipping at the same position (*i*=1), we define *S_i,j_* as the nucleotide at position *i* in clipped sequence *j*. We also define *T_i_* as the total number of overlapping clipped sequences at position *i*. We assume the first clipped position is *i*=1, and the last position with at least two clipped reads (*T_i_* > 1) is *i*=*S*. We first compute the proportion of each nucleotide *m* ∈ (A, C, T, G) at each position *i* as:

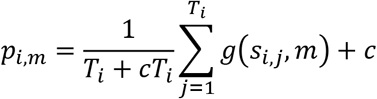

where *c* = 0.1 is a small pseudocount and *g* returns 1 if two nucleotides are equal:

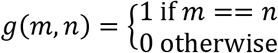

The Shannon entropy of a set of overlapping clipped sequences is then:

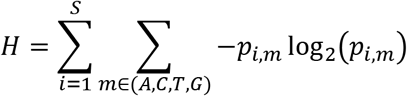

We define the information content of a set of clipped sequences as the difference between the observed entropy and the maximum possible entropy:

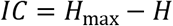

The maximum possible entropy, *H*_max_, is computed assuming that the nucleotides are evenly distributed across the four possible nucleotides at each site, taking into account that the number of sequences may not be divisible by 4. For example, if there are 6 clipped reads overlapping a position, the maximum possible entropy occurs when the four nucleotide proportions are 2/6, 2/6, 1/6, and 1/6.

### Training and testing BreakCA

BreakCA uses a random forest to predict whether a given 20bp window contains an indel using 16 features described in Supplementary Table 2. We created training and test datasets for the GM12878 lymphoblastoid cell line using ATAC-seq (50bp paired-end reads) data[13], H3K27ac ChIP-seq (50bp single-end reads) data[14] and indel genotypes from the Platinum Genomes Project[15]. We labeled 20bp windows centered around the start and end positions of indels within ChIP-seq or ATAC-seq peaks as “true” windows and 20bp windows located within peaks and not co-localizing with indels as “false” windows. We used 50% of the windows to create a training dataset and the remaining to create a test dataset. In total there were 2980 true (indel-containing) and 842,739 false (non indel-containing) windows in the training dataset and 2980 true and 842,738 false windows in the test dataset for paired-end ATAC-seq. For single-end ChIP-seq, there were 623 true and 254,328 false windows in the training dataset and 623 true and 254,327 false windows in the test dataset.

To implement the random forest model, we used the mlr package in R and tuned three hyperparameters by performing a grid search of reasonable hyperparameter values and choosing the values that yielded the highest accuracy (defined as mean (response == truth)) in 5-fold cross-validation. The 3 hyperparameters were ntree (number of trees to grow), mtry (number of predictors to use for node-split) and node-size (number of observations in the terminal node which is associated with the depth of the decision trees). After choosing hyperparameter values, we trained the random forest on the complete training dataset and applied it to the test dataset, using the fraction of true votes from the decision trees as the prediction score.

We compared the performance of BreakCA to two popular variant callers: VarScan2[16] and the GATK-HaplotypeCaller[17]. For VarScan2 and the GATK-HaplotypeCaller we extracted the start and end position of the indels using VariantAnnotation[49] and overlapped them with true and false windows in the test dataset. For VarScan2, we used 1.0 -*p-value* as the score and assigned the highest score to each window. For the GATK-HaplotypeCaller we used QD (phred-scaled variant call confidence normalized by allele depth) as the score and assigned the highest score to overlapping windows. For true and false windows with insufficient coverage to call variants, we set the prediction output to 0 for BreakCA, VarScan2, and GATK. To draw precision-recall curves and compute the area under them, we interpolated between datapoints by setting the precision at each recall level, *r*, to *p_interp_*(*r*) = max(*p*(*r*′)), where *r*′ ≥ *r* (see https://nlp.stanford.edu/IR-book/html/htmledition/evaluation-of-ranked-retrieval-results-1.html). We did not evaluate the performance of popular indel callers MANTA[50], pindel[51], or Delly[52] because all of these methods require paired-end reads and/or rely on the distribution of insert sizes from mapped reads, which limits the number of ChIP-seq and ATAC-seq datasets these callers can be applied to. In addition, the use of insert sizes is unlikely to work well for ATAC-seq reads, which have a multimodal distribution that changes depending on whether the reads are from nucleosome-free or nucleosome-containing genomic regions[13].

We further evaluated the performance of BreakCA using ATAC-seq from the Jurkat T-ALL cancer cell line. For this cell line we used indel genotypes from the GATK-HaplotypeCaller applied to WGS as “ground truth”[18]. In total, there were 5020 true and 1,023,159 false windows in the Jurkat ATAC-seq dataset and 6403 true and 1,267,161 false windows for the Jurkat H3K27ac ChIP-seq dataset. For these windows, we made predictions using the model trained on the GM12878 ATAC-seq training set and evaluated the precision-recall as described above.

### Overlapping variant windows with repeat regions

Genomic positions of known repeat sequences in the hg19 genome were downloaded from the UCSC Genome Browser’s RepeatMasker track. Short Tandem Repeat (STR) locations were obtained from Willems et al. 2017[53]. We used GenomicRanges (version 1.24.3)[54] to overlap indel windows with known repeat positions in the human genome and removed indel windows which overlapped STRs.

### Frequency of indels in gnomAD and GTEx

To filter indels that are likely to be common germline variants we overlapped our 20bp variant windows with indel calls from two large datasets: (1) 15,708 whole genomes from v2.1 of the genome aggregation database (gnomAD)[55] and (2) 635 whole genomes from V7 of the genotype-tissue expression project (GTEx)[29]. We used GATK-SelectVariants to identify gnomAD and GTEx indels overlapping our 20bp variant windows and used SNPSift to extract genomic position, reference and alternate allele and allele frequency (AF) fields from the VCF file[56]. We also required indels to have high coverage across gnomAD samples, removing those with a median coverage of less than 20 reads. For our analysis of rare germline/somatic indels, we retained variant windows with MAF ≤ 5× 10^−3^.

### Running BreakCA on GTEx samples

In addition to filtering based on indels identified by the gnomAD and GTEx variant calling pipelines, we added a filter for indels identified by BreakCA on 300 GTEx WGS samples[29]. This purpose of this filter is to remove common germline variants (or artefacts) that are detected by BreakCA, but that are not detected by GATK. We ran BreakCA on 300 GTEX WGS samples, using the same *α* and *β* prior distribution hyperparameters values that we estimated from the GM12878 ATAC-seq dataset. After running BreakCA, we calculated the fraction of samples (SF) with a predicted indel in each window (only considering samples with ≥10 reads overlapping a window as ‘testable’). (We use SF rather than MAF here because BreakCA calls indels as present/absent and does not distinguish between heterozygotes and homozygotes). For our analysis of rare germline/somatic indels, we retained variant windows with SF ≤ 5×10^−3^.

### Estimating the expected frequency of recurrent indels

To create Fig 3C, we calculate the expected number of windows, *T_x_*, that would contain RS indels in *x* cell lines, conditional on seeing an indel in one cell line:

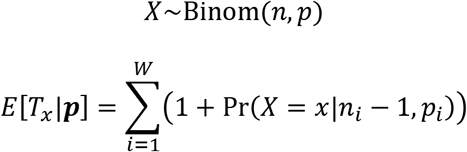

where *W* is the total number of windows, and ***p*** is a vector of length *W*, with elements *p_i_* that give the expected proportion of cells with an indel. We define *n_i_* is the total number of testable cell lines for window *i* (i.e. those with sufficient read depth), and subtract 1 from *n_i_* to account for the fact that we have conditioned on seeing an indel in one cell line already. We assume Hardy-Weinberg equilibrium and set 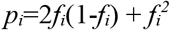, where *f_i_* is the allele frequency of the indel in window *i*. We set the allele frequency to either *f_i_*=2×10^−3^ for all windows, *f_i_*=1×10^−3^ for all windows, or *f_i_*=max(*gi*, 1×10^−4^), where *g_i_* is the observed allele frequency of the indel in gnomAD. For windows that contained multiple gnomAD indels, we used the one with the highest allele frequency. We use *f_i_*=1×10^−3^, because this is the gnomAD allele frequency cutoff used to identify RS indels. We use *f_i_*=2× 10^−3^ as a conservative assumption that gnomAD underestimates some allele frequencies. We use *f_i_*=max(*gi*, 1×10^−4^) to match observed allele frequencies in gnomAD (which are typically much lower than 1×10^−3^).

To calculate a p-value for the number of observed windows with RS indels in 4 or more cell lines we calculate the probability of observing 4 or more indels using the Poisson cumulative distribution function:

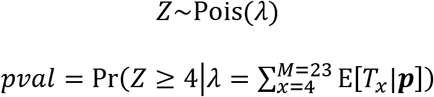

Where *λ* is the expected rate of windows containing 4 or more RS indels and M=23 is the number of cell lines in our study.

RS indels could also be observed due to elevated mutation rates within some windows or due to false positive indel calls. The former possibility is only likely if the window mutation rate is substantially exceeds the allele frequencies that we assume above. Even if we assume an allele frequency of 0.05% (equivalent to a window indel mutation rate of approximately 2p=0.001), the expected number of windows with 4 or more RS indels is 0.38, far lower than the observed 4 (p=6.4×10^−4^ by Poisson test). This number of windows with RS indels is also unlikely to result from indel call errors (assuming the errors occur independently) because we estimate the per-window false discovery rate for BreakCA on ChIP-seq data to be 1.7×10^−4^—an order of magnitude lower than the allele frequencies we assume above.

### Testing recurrent indels for association with gene expression

We used GenomicRanges (version 1.34.0) to find promoters of genes located within 100kb of recurrent rare germline or somatic (RS) indels and used Student’s t-test to test for differences in mean expression between the indel and non-indel groups. To test if the association with expression is expected by chance we permuted the sample labels for each test and compared the signals with quantile-quantile plots.

### Transcription factor motif discovery

We obtained the reference sequence (hg19/GRCh37) for 40bp regions centered around predicted indels and introduced the indel to create a non-reference sequence. We used TFBSTools[57] to search the JASPAR2016 database[58] for known transcription factor binding motifs located within the reference and the non-reference sequences. We filtered the motifs using p-value ≤ 0.001 (computed by TFMPvalue[59]) and kept only the top 10% of the motifs found uniquely in either reference or non-reference sequences as our most-reliable hits.

## Data and source code availability

ATAC-seq data from the Jurkat, RPMI-8402, MOLT-4, CCRF-CEM and K-562 cell lines has been submitted to GEO under accession GSE129086. The BreakCA source code is available from https://github.com/Arkosen/BreakCA.

## Author Contributions

GM and AS conceived of the project. GM supervised the project. AS developed the BreakCA software and analyzed the data. STT performed the ATAC-seq experiments on the T-ALL cell-lines. YF performed bioinformatic processing and QC of the ATAC-seq data under the supervision of GE. GM and AS wrote and edited the manuscript.

## Acknowledgements

This research was supported by NIH-NCI CCSG: P30 014195, and a grant from Padres Pedal the Cause/RADY #PTC2017. AS was supported by a Pioneer Fund Postdoctoral Scholar Award. Sequencing was carried out by the NGS Core Facility of the Salk Institute with funding from NIH-NCI CCSG: P30 014195, the Chapman Foundation, and the Helmsley Charitable Trust. We thank N. Hah and T. Nguyen for technical support. The Razavi Newman Integrative Genomics and Bioinformatics Core Facility of the Salk Institute is funded through the NIH-NCI CCSG: P30 014195 and the Helmsley Charitable Trust.

## Supplementary Figures

**Supplementary Figure 1:**
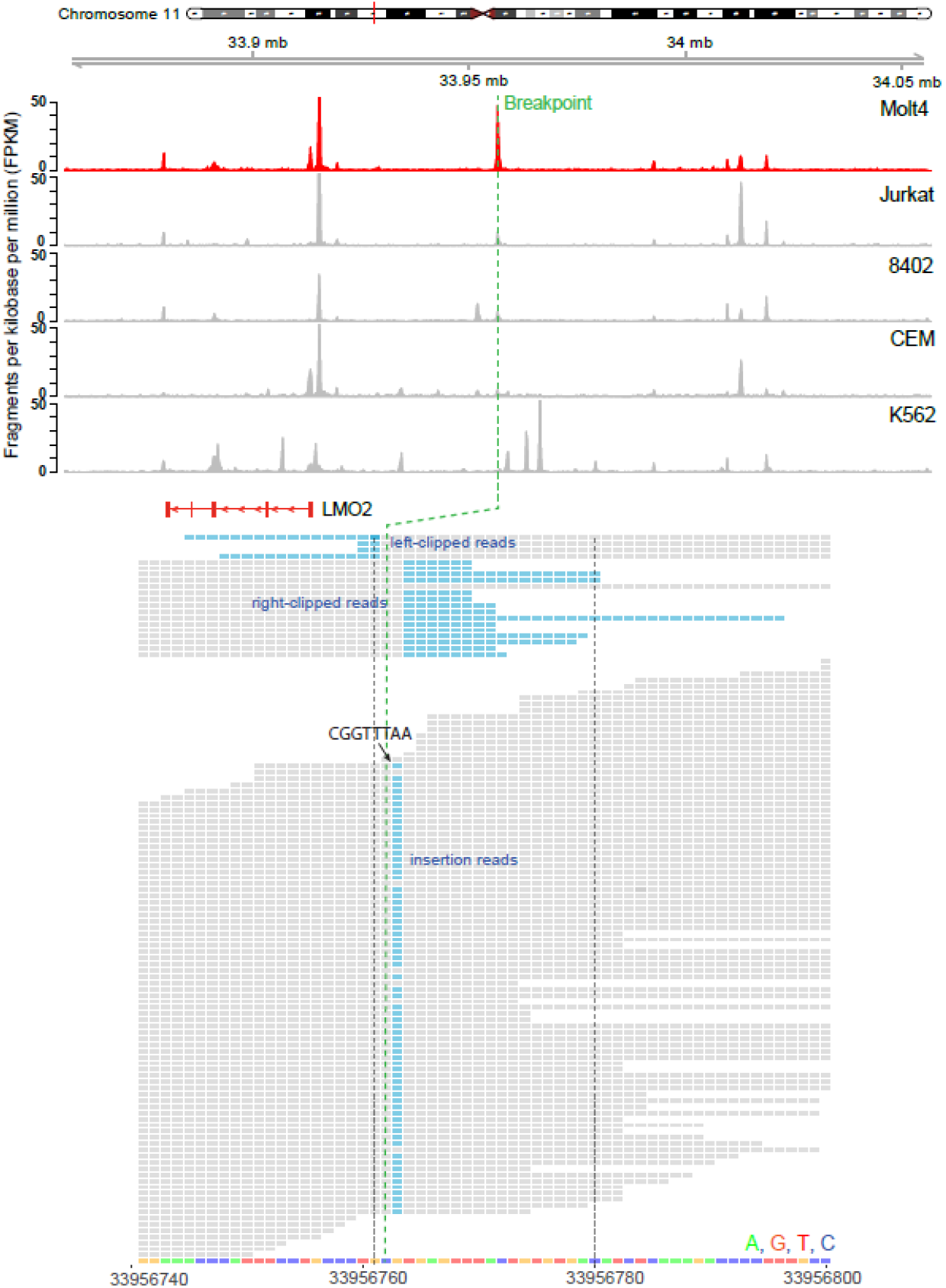
BreakCA detects a known 8bp CGGTTTAA insertion upstream of the LMO2 oncogene in the MOLT-4 cell line. Mapped read alignments are grey with soft-clipped bases and insertion positions indicated in blue.

**Supplementary Figure 2:**
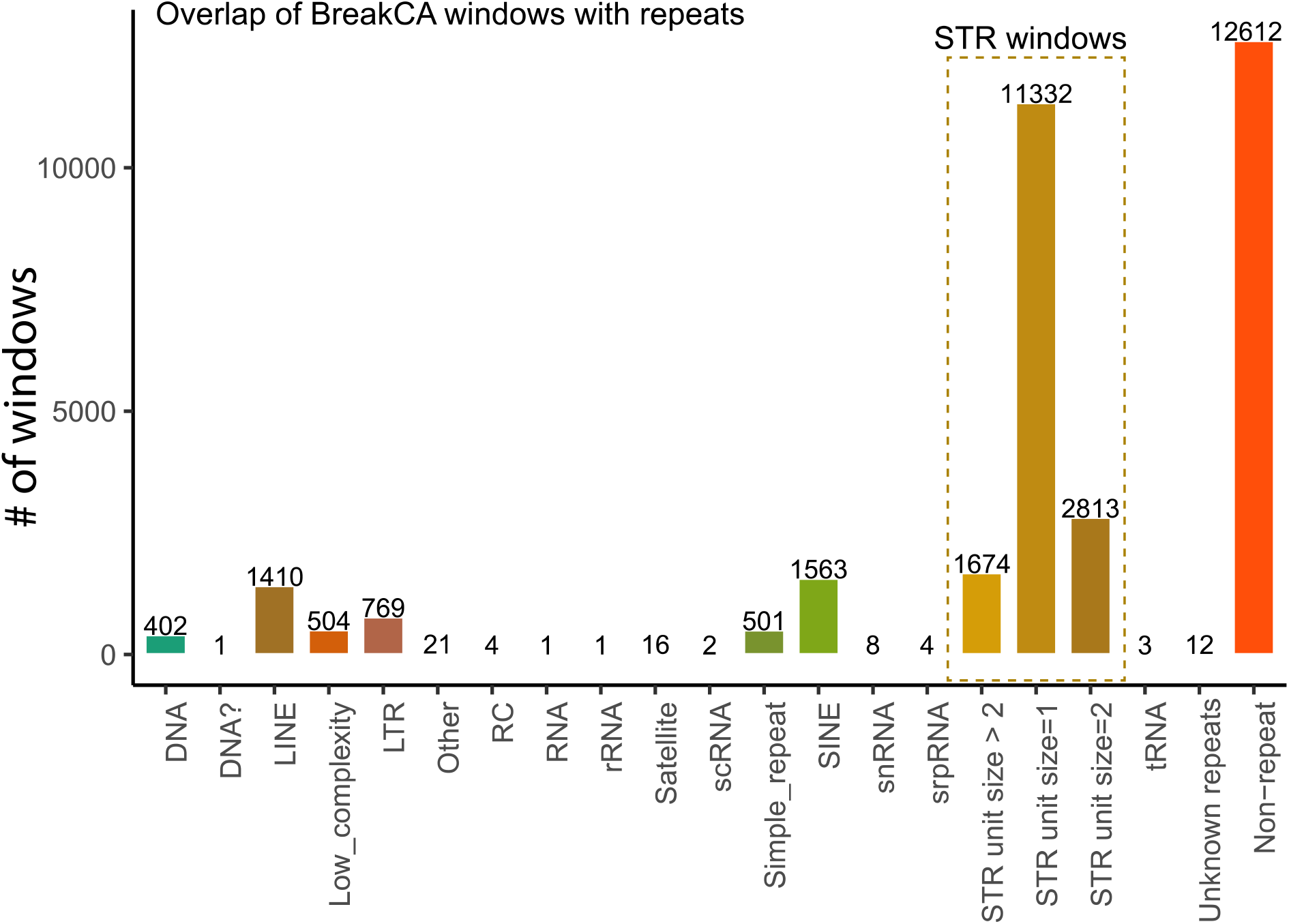
Overlap of indel windows with repeats regions from RepeatMasker or short tandem repeats (STRs). Indel windows overlapping STRs are not included in our analyses.

**Supplementary Figure 3:**
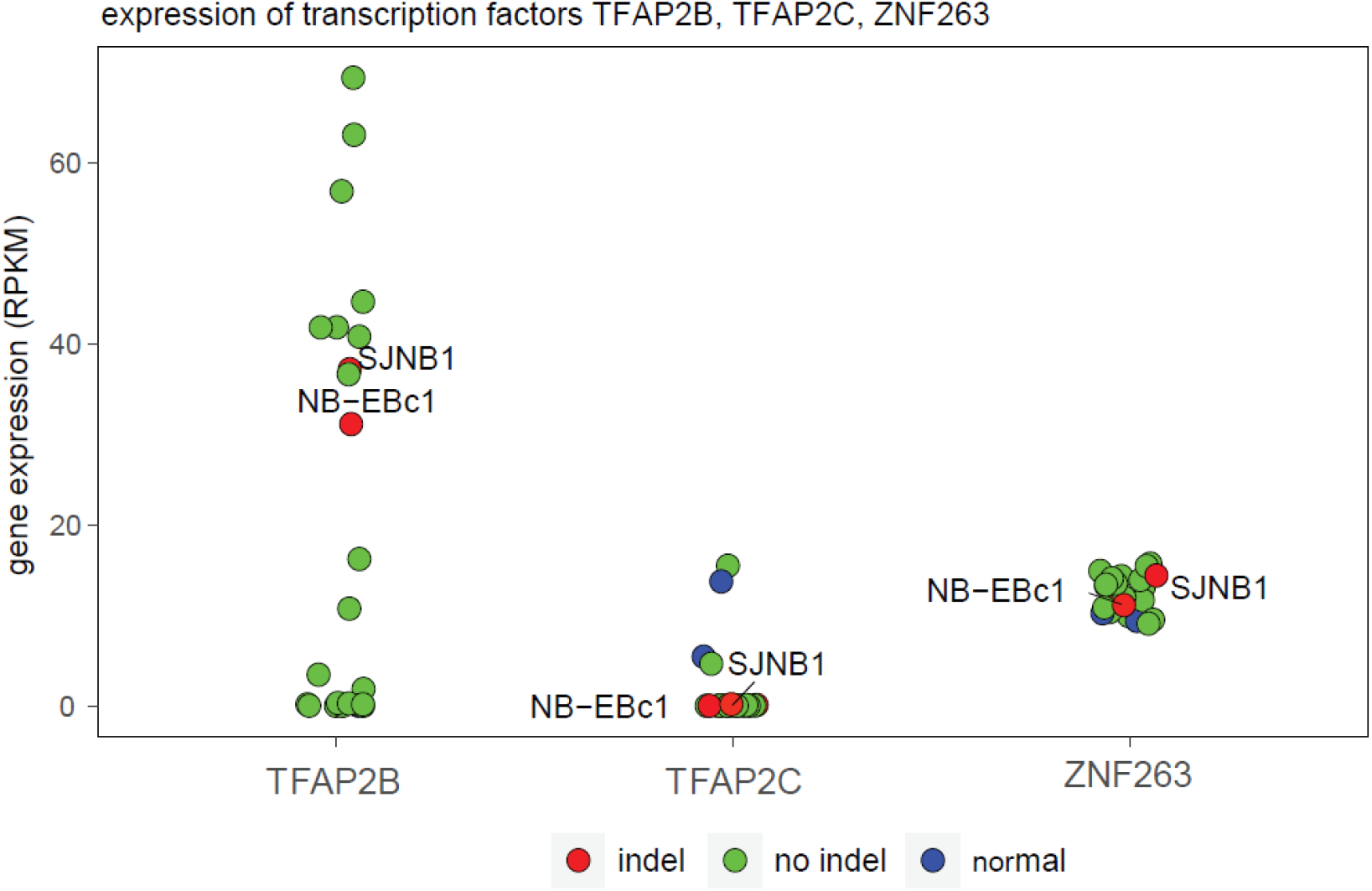
Gene expression of the *TFAP2B, TFAP2C* and *ZNF263* transcription factors in neuroblastoma cell lines and human neural crest cells (hNCCs) demonstrating that *TFAP2B* and *ZNF263* are expressed in the SJNB1 and NB-EBc1 cell lines, while *TFAP2C* is not expressed in NB cell-lines.

**Supplementary Figure 4:**
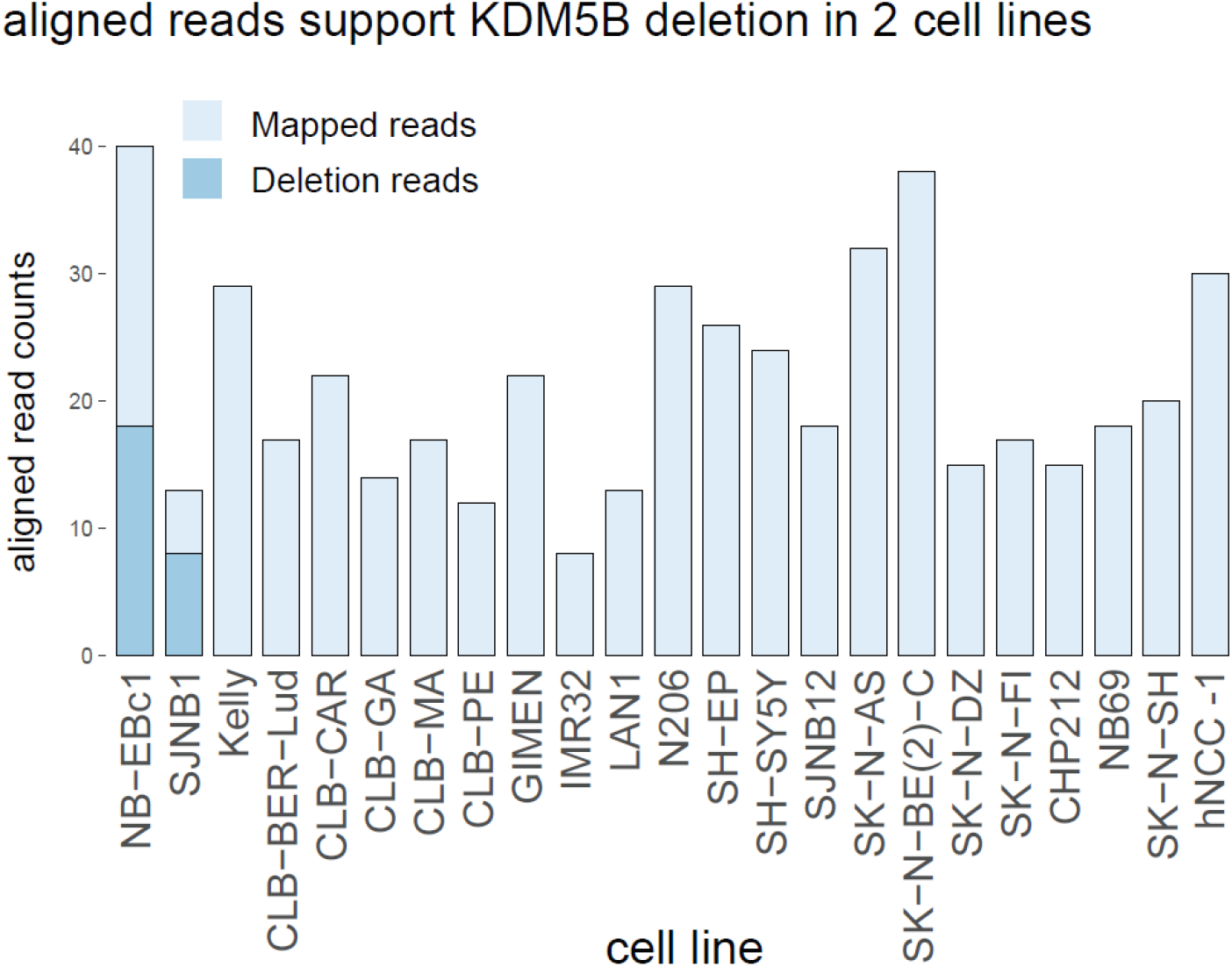
The count of aligned reads that contain deletions at position chr1:202777149. Only the SJNB1 and NB-EBc1 cell lines have any deletion reads at this position.

**Supplementary Figure 5:**
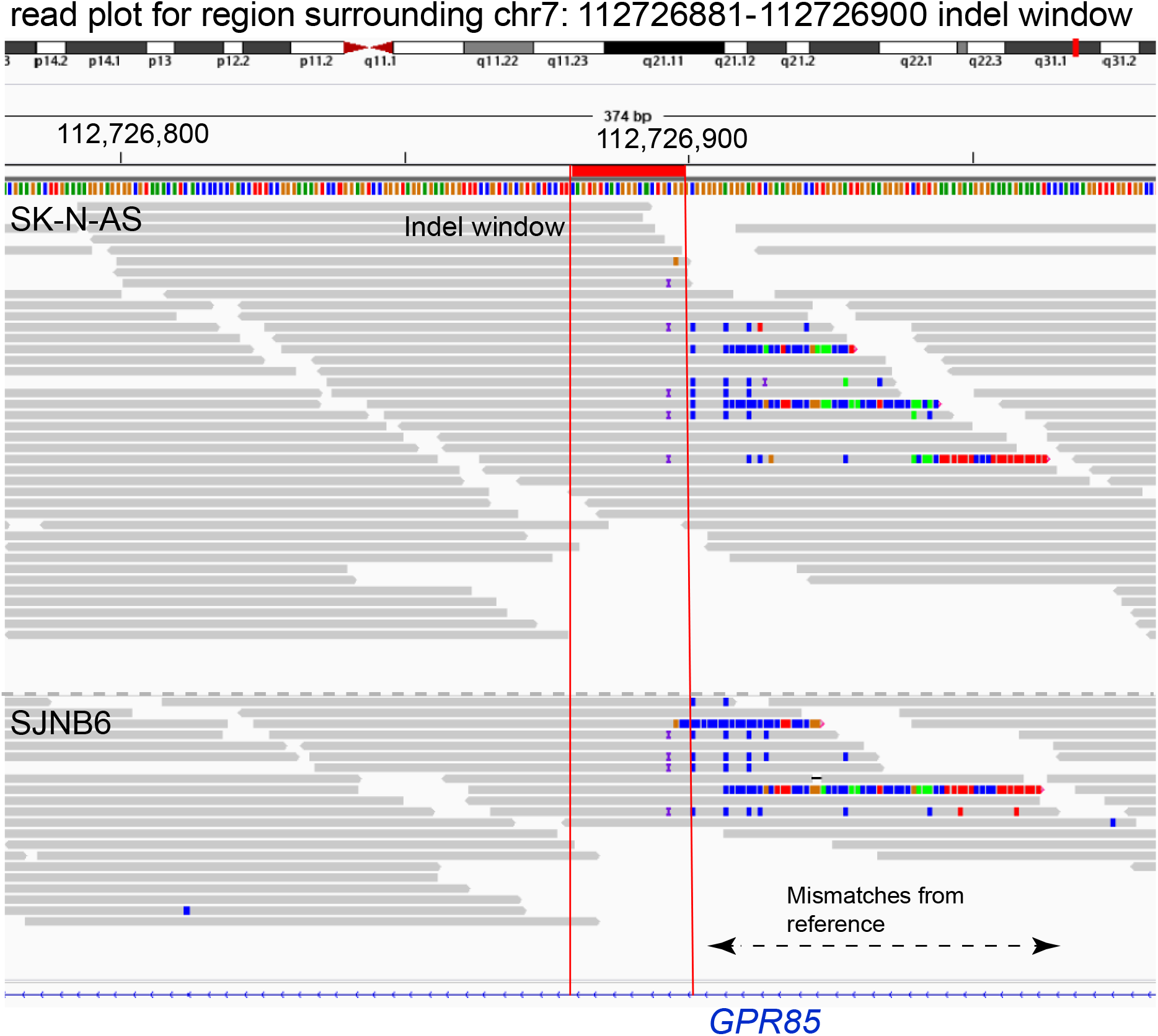
Read plots from the Integrative Genomics Viewer[60] for two of the neuroblastoma cell lines containing the complex event in the GPR85 intron. Reads are colored when they are clipped or have a base mismatch from the reference. Insertions are indicated with a small purple “I”.

## Supplementary Tables

**Supplementary Table 1:** Read statistics including the fraction of reads in peaks (FRiP) for ATAC-seq datasets generated from the Jurkat, CCRF-CEM, RPMI-8402, MOLT-4, and K-562 cell lines.

**Supplementary Table 2:** Description of prediction features used by the BreakCA random forest.

**Supplementary Table 3:** List of rare-germline and somatic indels detected in 23 NB cell-lines with features used for BreakCA prediction.

